# The complete mitochondrial genome sequence of *Oryctes rhinoceros* (Coleoptera: Scarabaeidae) based on long-read nanopore sequencing

**DOI:** 10.1101/2020.04.27.065235

**Authors:** Igor Filipović, James P. Hereward, Gordana Rašić, Gregor J. Devine, Michael J. Furlong, Kayvan Etebari

## Abstract

The coconut rhinoceros beetle (CRB, *Oryctes rhinoceros*) is a severe and invasive pest of coconut and other palms throughout Asia and the Pacific. The biocontrol agent, Oryctes rhinoceros nudivirus (OrNV), has successfully suppressed *O. rhinoceros* populations for decades but new CRB invasions started appearing after 2007. A single-SNP variant within the mitochondrial *cox1* gene is used to distinguish the recently-invading CRB-G lineage from other haplotypes, but the lack of mitogenome sequence for this species hinders further development of a molecular toolset for biosecurity and management programmes against CRB. Here we report the complete circular sequence and annotation for CRB mitogenome, generated to support such efforts.

Sequencing data were generated using long-read Nanopore technology from genomic DNA isolated from a CRB-G female. The mitochondrial genome was assembled with Flye v.2.5, using the short-read Illumina sequences to remove homopolymers with Pilon, and annotated with MITOS. Independently-generated transcriptome data were used to assess the *O. rhinoceros* mitogenome annotation and transcription. The aligned sequences of 13 protein-coding genes (PCGs) (with degenerate third codon position) from *O. rhinoceros*, 13 other Scarabaeidae taxa and two outgroup taxa were used for the phylogenetic reconstruction with the Maximum likelihood (ML) approach in IQ-TREE and Bayesian (BI) approach in MrBayes.

The complete circular mitochondrial genome of *O. rhinoceros* is 20,898 bp-long, with a gene content canonical for insects (13 PCGs, 2 rRNA genes, and 22 tRNA genes), as well as one structural variation (rearrangement of *trnQ* and *trnI*) and a long control region (6,204 bp). Transcription was detected across all 37 genes, and interestingly, within three domains in the control region. ML and BI phylogenies had the same topology, correctly grouping *O. rhinoceros* with one other Dynastinae taxon, and recovering the previously reported relationship among lineages in the Scarabaeidae. *In silico* PCR-RFLP analysis recovered the correct fragment set that is diagnostic for the CRB-G haplogroup. These results validate the high-quality of the CRB mitogenome sequence and annotation.

## Introduction

*Oryctes rhinoceros* (Linnaeus 1758) (Coleoptera: Scarabeidae: Dynastinae), also known as the coconut rhinoceros beetle (CRB), is an important agricultural pest causing significant economic damage to coconut and other palms across Asia and South Pacific. During the 20th century human mediated dispersal resulted in the distribution of *O. rhinoceros* expanding from its native range (between Pakistan and the Philippines) throughout Oceania (Catley 1969). After the discovery and introduction of the viral biocontrol agent Oryctes rhinoceros nudivirus (OrNV) in the 1960s, most of the CRB populations in the Pacific islands have been persistently suppressed (Huger 2005). However, after a biocontrol campaign failed to eradicate a newly established population in Guam in 2007, new CRB invasions were recorded in Papua New Guinea (2009), Hawaii (2013) and Solomon Islands (2015) (Marshall et al. 2017). Worryingly, the new invasive populations have also been difficult to control by known OrNV isolates (Marshall et al. 2017), emphasizing the importance of actively overseeing and adapting the management programmes for this important insect pest.

The expansion pathways, dynamics and hybridization of invasive insect pests and other arthropods are commonly traced through the analyses of mitochondrial sequence variation (e.g. (Wang et al. 2017; Rubinoff et al. 2010; Moore et al. 2013)). In the absence of a mitochondrial genome sequence, the universal barcoding region is often amplified with degenerate primers to investigate partial sequences of *cox1* and a limited number of other mitochondrial genes in the target species. However, analyses of such partial sequence data can fail to distinguish true mitochondrial lineages unless a sufficient number of genetic markers can be retrieved. Variation from a partial sequence of one mitochondrial gene (*cox1*) and one nuclear gene (*cad*) was not sufficient to allow confident hypotheses testing around *O. rhinoceros* invasion pathways (Reil et al. 2016), but a single diagnostic SNP within the partial *cox1* gene amplicon has been used to distinguish the CRB-G haplotype from other haplotype that originally invaded the Pacific islands in the early 1900s (Marshall et al. 2017). Here we report the first and complete mitochondrial genome sequence assembly of *O. rhinoceros*, a genomic resource that will support the development of a comprehensive molecular marker toolset to help advance the biosecurity and management efforts against this resurgent pest.

The complete *O. rhinoceros* mitogenome assembly was generated using long-read Oxford Nanopore Technologies (ONT) sequencing and complemented with the short-read Illumina sequencing. The approach recovered all genes in the canonical order for insects and a long non-coding (control) region (6,204 bp) that was absent from a short-read assembly, probably because it contained different putative tandem repeats. Three spots with detectable transcription within control region and the rearrangement of two tRNA genes (*trnI and trnQ*) were also identified. The high quality of the assembly was validated through the correct placement of *O. rhinoceros* within the Scarabaeidae phylogeny, transcription patterns from an independently-generated transcriptome dataset, and *in silico* recovery of a recently reported diagnostic PCR-RFLP marker. This is the first complete mitogenome for the genus *Oryctes* and the subfamily Dynastinae, and among only a few for the entire scarab beetle family (Scarabaeidae).

## Materials and Methods

### Sample collection, DNA extraction and ONT sequencing

An adult female *O. rhinoceros* female was collected from a pheromone trap (Oryctalure, P046-Lure, ChemTica Internacional, S. A., Heredia Costa Rica) on Guadalcanal, Solomon Islands in January 2019 and preserved in 95% ethanol. Initially, the mitochondrial haplotype of the specimen was established as CRB-G (Marshall et al. 2017) *via* Sanger sequencing of the partial *cox1* gene sequence that was amplified using the universal barcode primers LCO1490 and HCO2198 (Folmer et al. 1994). High-molecular weight DNA was extracted using a customized magnetic (SPRI) bead-based protocol. Specifically, smaller pieces of tissue from four legs and thorax (50 mm3) were each incubated in a 1.7 ml eppendorf tube with 360 μL ATL buffer, 40 μL of proteinase K (Qiagen Blood and Tissue DNA extraction kit) for 3h at RT, while rotating end-over-end at 1 rpm. 400 μL of AL buffer was added and the reaction was incubated for 10 min, followed by adding 8 μL of RNase A and incubation for 5 minutes. Tissue debris was spun down quickly (1 min at 16,000 rcf) and 600 μL of homogenate was transferred to a fresh tube, where SPRI bead solution was added in 1:1 ratio and incubated for 30 min while rotating at end-over-end at 1 rpm. After two washes with 75% ethanol, DNA was eluted in 50 μL of TE buffer. DNA quality (integrity and concentration) was assessed on the 4200 Tapestation system (Agilent) and with the Qubit broad-range DNA kit. To enrich for DNA >10 kb, size selection was done using the Circulomics Short Read Eliminator XS kit. We sequenced a total of four libraries, each prepared with 1 μg of size-selected HMW DNA, following the manufacturer’s guidelines for the Ligation Sequencing Kit SQK-LSK109 (Oxford Nanopore Technologies, Cambridge UK). Sequencing was done on the MinION sequencing device with the Flow Cell model R9.4.1 (Oxford Nanopore Technologies) and the ONT MinKNOW Software.

An Illumina sequencing library was also prepared using a NebNext Ultra DNA II Kit (New England Biolabs, USA). The library was sequenced on a HiSeq X10 (150bp paired end reads) by Novogene (Beijing, China).

### Genome assembly, annotation and analysis

The Guppy base caller ONT v.3.2.4 was used for high-accuracy base calling on the raw sequence data, and only high-quality sequences with a Phred score > 13 were used for the *de novo* genome assembly with the program Flye v.2.5 (Kolmogorov et al. 2019) in the metagenome assembly mode. The method recovered the full circular assembly and to verify its accuracy, we first mapped the original reads back to the generated mitogenome assembly using Minimap2 (Heng Li 2018) with the following parameters: -k15 --secondary=no -L −2. Second, we used BWA-MEM (H. Li and Durbin 2009) to map short-read Illumina sequences obtained from the whole-genome sequencing of another *O. rhinoceros* female collected from the same geographic location as the sample used for the mitogenome assembly. The read alignment analysis in Pilon (Walker et al. 2014) was used to identify inconsistencies between the draft mitogenome assembly and the aligned short Illumina reads, removing small indels that represent homopolymers (e.g. >4bp single nucleotide stretches) as an inherent sequencing error of the ONT (Mikheyev and Tin 2014). Finally, we manually inspected if the Pilon correction occurred only in putative homopolymer regions by comparing the draft assembly with the Pilon-polished version.

The complete mitogenome sequence was initially annotated using the MITOS web server (Bernt et al. 2013), and tRNA genes and their secondary structures were cross-analysed using tRNAscan-SE v2.0 (Chan et al., 2019). To further refine the annotation and to examine mitogenome transcription, we used BWA-MEM to align Illumina reads from a transcriptome study of *O. rhinoceros* larvae (Shelomi, Lin, and Liu 2019), retrieved from the NCBI (SRR9208133). Finally, we manually inspected and compared our annotation to the complete and near complete mitogenome annotations of other related taxa (Table 1) in Geneious (2020.0.4). MEGA X (Kumar et al. 2018) was used to assess the codon usage and nucleotide composition of protein-coding genes. We used Geneious (2020.0.4) to test if the nucleotide sequence of the *cox1* gene recovers the recently reported PCR-RFLS marker (Marshall et al. 2017). This was done by aligning the sequences of the primer pair (LCO1490 and HCO2198) to isolate the amplicon fragment and by performing *in silico* restriction digestion with MseI restriction enzyme. The restriction digestion of the amplicon produces a set of fragment lengths that distinguishes CRB-G from other haplotypes (Marshall et al. 2017). The presence of tandem repeats within the control region was assessed with the Tandem Repeats Finder v.4.0.9 (Benson 1999) using default parameters. The annotated mitochondrial DNA (mtDNA) sequence has been deposited in GenBank under accession number (will be available later).

**Table 1.** Taxa with complete or partial metagenome sequences used for the phylogenetic analyses.

| Accession | Organism | Genome type | Missing genes | Contains control region | Sequence Length (bp) |
| --- | --- | --- | --- | --- | --- |
| FJ859903.1 | *Rhopaea magnicornis* | complete | none | yes | 17522 |
| JX412731.1 | *Cyphonistes vallatus* | partial | nad1 | no | 11629 |
| JX412734.1 | *Trox* sp. TRO01 | partial | nad2; cox1 | no | 11622 |
| JX412739.1 | *Schizonycha* sp. SCH01 | partial | nad2 | no | 13542 |
| JX412755.1 | *Asthenopholis* sp. AST01 | partial | nad2 | no | 12352 |
| KC775706.1 | *Protaetia brevitarsis* | complete | none | yes | 20319 |
| KF544959.1 | *Polyphylla laticollis mandshurica* | partial | none | no | 14473 |
| KU739455.1 | *Eurysternus foedus* | partial | none | no | 15366 |
| KU739465.1 | *Coprophanaeus* sp. BMNH679884 | partial | none | no | 15554 |
| KU739469.1 | *Bubas bubalus* | partial | none | no | 16035 |
| KU739498.1 | *Onthophagus rhinolophus* | partial | none | no | 15237 |
| KX087316.1 | *Melolontha hippocastani* | partial | none | no | 15485 |
| MN122896.1 | *Anoplotrupes stercorosus* | partial | nad2 | no | 13745 |
| NC_030778.1 | *Osmoderma opicum* | complete | none | yes | 15341 |
| NC_038115.1 | *Popillia japonica* | complete | none | yes | 16541 |

### Phylogenetic analysis

To ascertain if our newly sequenced CRB mitogenome can be correctly placed within the Dynastinae subfamily of the Scarabaeidae family, we performed the phylogenetic analyses with 15 additional taxa for which complete or near complete mitogenome sequences were available in NCBI. We used thirteen species from five subfamilies of the Scarabaeidae family (Dynastinae, Rutelinae, Cetoniine, Melolonthinae, Scarabaeinae) and members of two other families from Scarabaeoidea as outgroups (Trogidae, Geotrupidae) (Table 1).

Nucleotide sequences of all 13 protein-coding genes (PCGs) were first translated into amino acid sequences under the invertebrate mitochondrial genetic code and aligned using the codon-based multiple alignment in Geneious (2020.0.4). The aligned amino acid matrix was back-translated into the corresponding nucleotide matrix and the Perl script Degen v1.4 (Zwick, Regier, and Zwickl 2012; Regier et al. 2010) was used to create the degenerated protein-coding sequences in order to reduce the bias effect of synonymous mutations on the phylogenetic analysis. These final alignments from all 13 PCGs were concatenated using Geneious (2020.0.4).

We estimated the phylogeny using two methods: the Maximum likelihood (ML) inference implemented in IQ-TREE web server (Trifinopoulos et al. 2016), and the Bayesian inference in MrBayes (Huelsenbeck and Ronquist 2001). For the ML analysis, the automatic and FreeRate heterogeneity options were set under optimal evolutionary models, and the branch support values were calculated using the ultrafast bootstrap (Hoang et al. 2018) and the SH-aLRT branch test approximation (Shimodaira and Hasegawa 1999) with 1000 replicates. For the BI analysis, we used the GTR+I+G substitution model, a burn-in of 100,000 trees and sampling every 200 cycles. The consensus trees with branch support were viewed and edited in Figtree v1.4.2.

## Results

### Mitogenome composition, organization and transcription

The ONT long reads enabled the complete assembly of the circular mitochondrial genome for *O. rhinoceros*, with a median coverage of >10,000x over the entire sequence length of 20,898 bp. This is the only complete mitogenome assembly and annotation for the Dynastinae subfamily, and among only a few complete mitogenomes for the scarab beetles. Our Illumina-only assembly recovered a 17,665bp mitogenome assembly, and this extra 3,233bp represents a repetitive control-region that the Illumina data will not map to. There are three transcriptionally active regions in this extra part of the mitogenome that only long reads have been able to reveal.

The annotation revealed a gene order canonical for insects (Cameron 2014), except for the rearrangement of *trnI* and *trnQ* genes, that showed the order: CR (control region)-*<u>trnQ-trnI</u>-trnM-nd2* instead of *CR-trnI-<u>trnQ-trnM</u>-nd2* (Table 2). The control region (CR) resided within a large non-protein coding region (6,204 bp long) located between *rrnS* and *trnQ*, and the Tandem Repeats Finder Analysis revealed a complex structure of this large region with 11 putative repeats that had a consensus sequence between 7 and 410 bp repeated 2-12 times (Table 3). The length of 22 tRNAs ranged from 63 to 70 bp (Table 2), and their predicted secondary structures exhibited a typical clover-leaf structure. The length of *rrnL* and *rrnS* were 1,283 bp and 783 bp, respectively (Table 2).

**Table 2.** Organization of the newly sequenced mitogenome of *Oryctes rhinoceros*.

| Feature name | Type | Start position | End position | Length | Direction | Start codon | Stop codon |
| --- | --- | --- | --- | --- | --- | --- | --- |
| trnQ | tRNA | 1 | 69 | 69 | reverse |  |  |
| trnI | tRNA | 127 | 190 | 64 | forward |  |  |
| trnM | tRNA | 195 | 263 | 69 | forward |  |  |
| nad2 | gene | 276 | 1271 | 996 | forward | ATT | TAA |
| trnW | tRNA | 1270 | 1335 | 66 | forward |  |  |
| trnC | tRNA | 1328 | 1392 | 65 | reverse |  |  |
| trnY | tRNA | 1393 | 1456 | 64 | reverse |  |  |
| cox1 | gene | 1458 | 2993 | 1536 | forward | ATC | TAA |
| trnL2 | tRNA | 2989 | 3054 | 66 | forward |  |  |
| cox2 | gene | 3055 | 3762 | 708 | forward | ATA | TAA |
| trnK | tRNA | 3743 | 3812 | 70 | forward |  |  |
| trnD | tRNA | 3813 | 3875 | 63 | forward |  |  |
| atp8 | gene | 3876 | 4031 | 156 | forward | ATT | TAA |
| atp6 | gene | 4025 | 4696 | 672 | forward | ATG | TAT |
| cox3 | gene | 4695 | 5483 | 789 | forward | ATG | TAT |
| trnG | tRNA | 5482 | 5545 | 64 | forward |  |  |
| nad3 | gene | 5546 | 5899 | 354 | forward | ATC | TAG |
| trnA | tRNA | 5898 | 5962 | 65 | forward |  |  |
| trnR | tRNA | 5963 | 6027 | 65 | forward |  |  |
| trnN | tRNA | 6028 | 6092 | 65 | forward |  |  |
| trnS1 | tRNA | 6093 | 6159 | 67 | forward |  |  |
| trnE | tRNA | 6161 | 6224 | 64 | forward |  |  |
| trnF | tRNA | 6223 | 6288 | 66 | reverse |  |  |
| nad5 | gene | 6287 | 8005 | 1719 | reverse | ATT | TAT |
| trnH | tRNA | 8003 | 8066 | 64 | reverse |  |  |
| nad4 | gene | 8066 | 9403 | 1338 | reverse | ATG | TAA |
| nad4l | gene | 9397 | 9687 | 291 | reverse | ATG | TAA |
| trnT | tRNA | 9690 | 9754 | 65 | forward |  |  |
| trnP | tRNA | 9755 | 9819 | 65 | reverse |  |  |
| nad6 | gene | 9821 | 10321 | 501 | forward | ATC | TAA |
| cob | gene | 10321 | 11463 | 1143 | forward | ATG | TAG |
| trnS2 | tRNA | 11462 | 11527 | 66 | forward |  |  |
| nad1 | gene | 11547 | 12497 | 951 | reverse | ATT | TAA |
| trnL1 | tRNA | 12499 | 12561 | 63 | reverse |  |  |
| rrnL | rRNA | 12559 | 13841 | 1283 | reverse |  |  |
| trnV | tRNA | 13843 | 13912 | 70 | reverse |  |  |
| rrnS | rRNA | 13912 | 14694 | 783 | reverse |  |  |
| Control region | misc_feature | 14695 | 20898 | 6204 | none |  |  |
| Expressed region 1 | misc_RNA | 15117 | 15253 | 137 | reverse |  |  |
| Expressed region 2 | misc_RNA | 15321 | 15476 | 156 | reverse |  |  |
| Expressed region 3 | misc_RNA | 17744 | 18387 | 644 | reverse |  |  |

**Table 3.** Characteristics of the putative tandem repeats in the control region of the *Oryctes rhinoceros* mitogenome.

| Indices | Period Size | Copy Number | Consensus Size | Percent Matches | Percent Indels | Score | A | C | G | T | Entropy (0-2) |
| --- | --- | --- | --- | --- | --- | --- | --- | --- | --- | --- | --- |
| 14764-14795 | 16 | 2 | 16 | 93 | 0 | 55 | 53 | 0 | 0 | 46 | 1 |
| 14757-14797 | 7 | 6 | 7 | 78 | 16 | 50 | 48 | 0 | 0 | 51 | 1 |
| 14818-14859 | 15 | 3 | 14 | 86 | 10 | 59 | 38 | 0 | 0 | 61 | 0.96 |
| 15012-15440 | 133 | 3.2 | 133 | 97 | 1 | 824 | 52 | 15 | 6 | 25 | 1.65 |
| 15447-17166 | 285 | 6 | 285 | 98 | 0 | 3336 | 24 | 29 | 13 | 33 | 1.93 |
| 16955-17949 | 206 | 4.9 | 205 | 98 | 1 | 1933 | 24 | 29 | 12 | 33 | 1.92 |
| 16955-17949 | 410 | 2.4 | 409 | 98 | 1 | 1938 | 24 | 29 | 12 | 33 | 1.92 |
| 18024-18419 | 110 | 3.6 | 110 | 96 | 1 | 724 | 41 | 24 | 8 | 25 | 1.83 |
| 18656-18707 | 22 | 2.3 | 23 | 81 | 15 | 65 | 51 | 1 | 3 | 42 | 1.31 |
| 19664-20895 | 102 | 12 | 102 | 98 | 0 | 2351 | 36 | 11 | 7 | 44 | 1.69 |
| 19664-20895 | 205 | 6 | 204 | 98 | 0 | 2369 | 36 | 11 | 7 | 44 | 1.69 |

The nucleotide composition of the CRB mitogenome sequence had high A + T bias (37.7% A, 32.8% T, 19.4% C and 10% G), which is highly concordant with other scarab beetle species, and the long CR matched this genome-wide pattern (34.3% A, 35.8% T, 20.4% C and 9.5% G).

All PCGs started with a standard initiation codon (ATN), ten of 13 PCGs terminated with the conventional stop codons (TAG or TAA), while three genes (*atp6, cox3* and *nad5*) had an incomplete stop codon T (Table 2). It is generally accepted that a cessation of protein translation can be signaled by incomplete codon structures in insects and other invertebrates (Cheng et al. 2016). *In silico* digestion of the *cox1* amplicon (delineated with the primer sequences from (Folmer et al. 1994)) produced the fragments 253 bp-,138 bp- and 92 bp-long (Supplemental Figure 1) that are diagnostic for the CRB-G haplotype (Marshall et al. 2017).

The mapping of the transcriptome sequencing reads to the newly assembled mitogenome revealed that all PCGs were transcribed (mean coverage depth per base >23,000, Figure 1), with *cox1* and *cox2* showing the highest level of expression when compared to the rRNA genes in the examined larval samples (Figure 1). We found three domains within the large CR-containing region that also showed detectable transcription levels, with the transcript sizes of 137, 156 and 644 bp respectively. Our attempts to annotate these transcripts were not successful due to the fact that their sequences did not contain any open reading frames, nor did they have any significant BLAST hits within the NCBI’s reference RNAseq or nucleotide databases.

**Figure 1.**
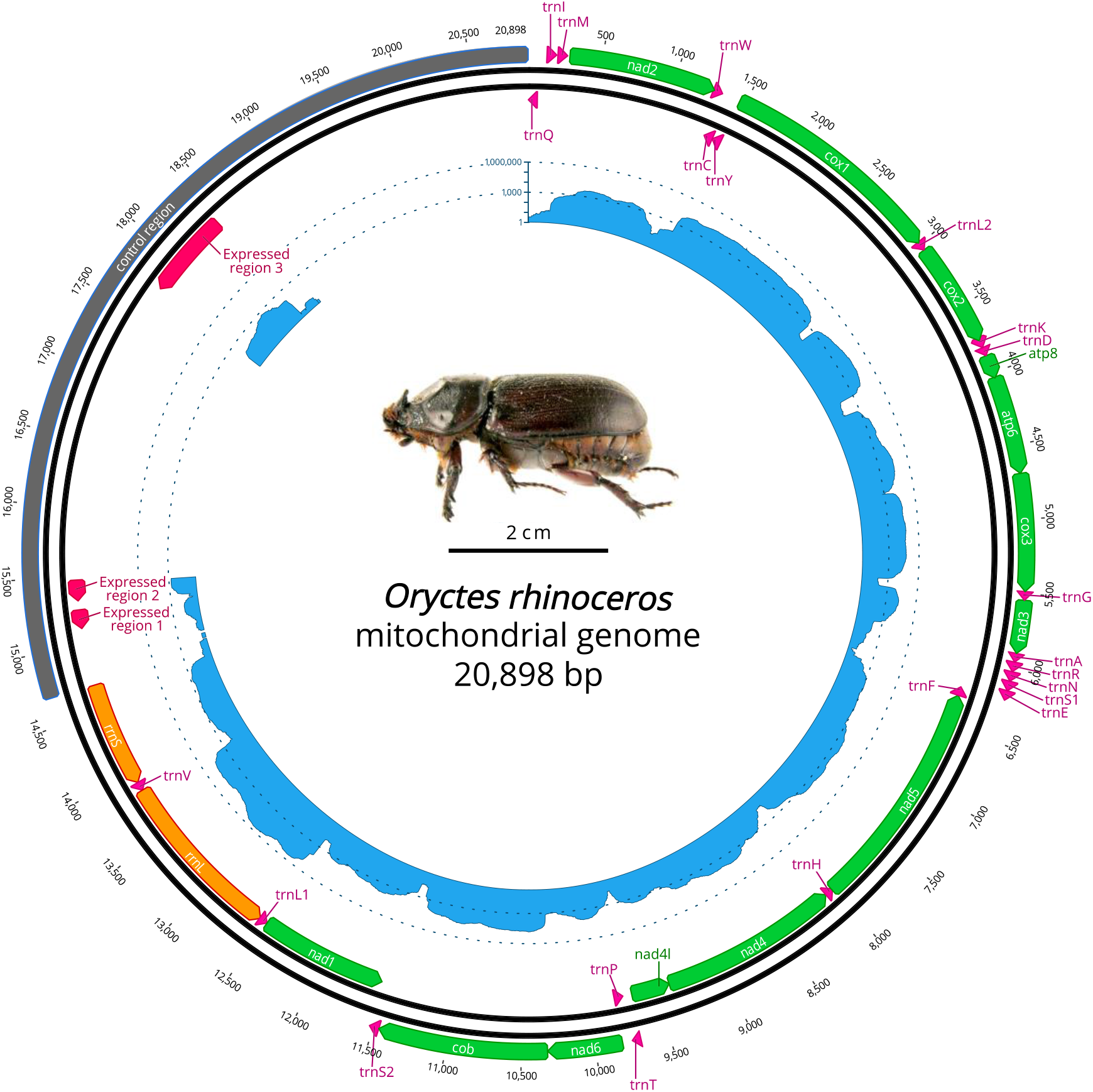
Circular representation of complete *O. rhinoceros* mitochondrial genome. The position and orientation of 13 PCG genes (green), 22 trna genes (pink), 2 rrna genes (orange), control region (grey) with 3 expressed domains (red). The inner circle displays transcriptome read depth (blue) on a logarithmic scale. Photo credit: *Oryctes rhinoceros* female, modified from Walker, K. (2005), Available online: PaDIL – http://www.padil.gov.au under the Creative Commons Attribution 3.0 Australia License.

### Phylogenetic analysis

Both ML and BI phylogenies grouped *O. rhinoceros* and another member of the Dynastinae subfamily (*Cyphonistes vallatus*) with 100% support (Figure 2, Supplemental Figure 2), and recovered the relationship of Dynastinae and Rutelinae as sister clades, that together with Cetoniine and Melolonthinae formed a basal split between phytophagus and coprophagus scarab beetles (Scarabaeinae) (Figure 2, Supplemental Figure 2). This phylogenetic reconstruction is highly congruent with the previously published phylogeny of scarab beetles (N. Song and Zhang 2018) and it groups *O. rhinoceros* with another member of the rhinoceros beetle subfamily (Dynastinae) with high confidence (100% SH-aLRT support, 100% ultrafast bootstrap support in ML, posterior probability 100% in BI), confirming the high quality of the newly described *O. rhinoceros* mitogenome.

**Figure 2.**
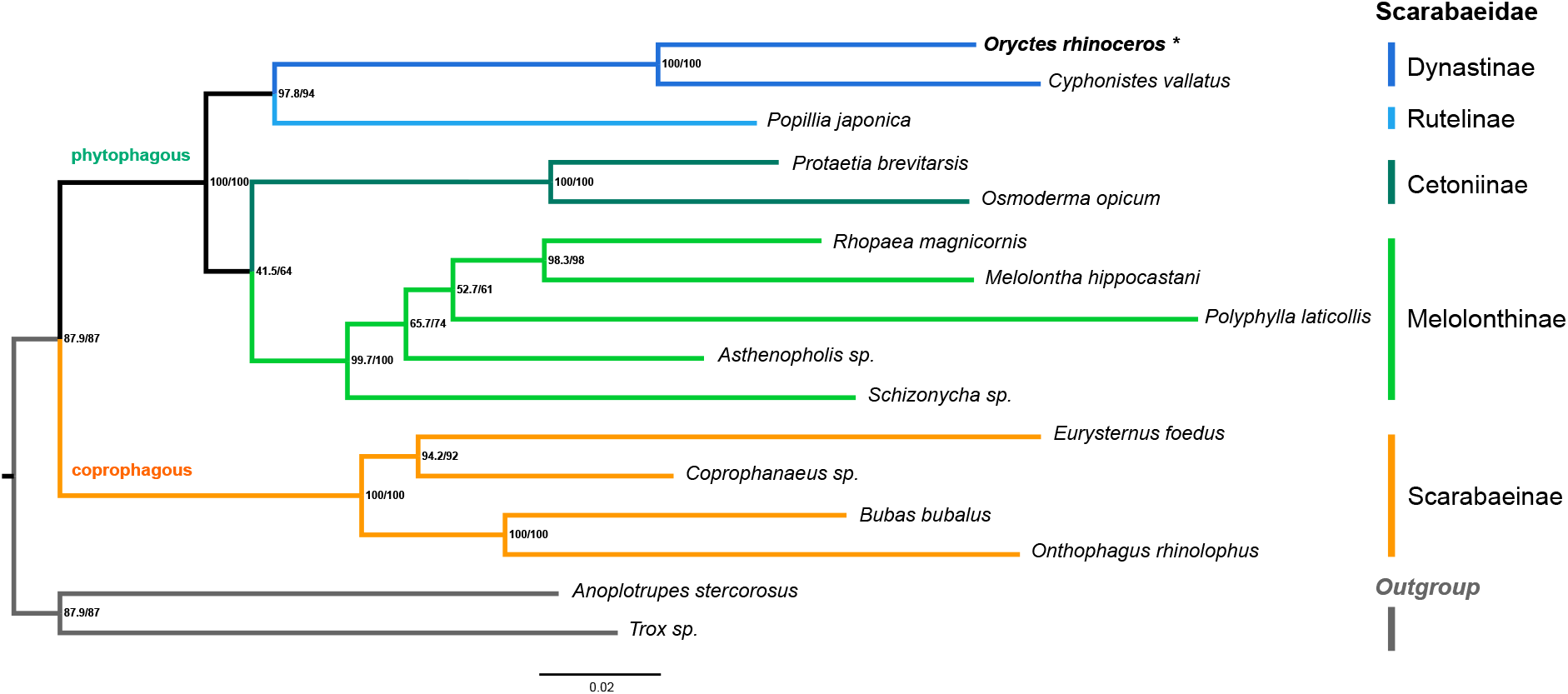
Maximum likelihood consensus tree inferred from the PCG dataset using IQ-TREE. Branch support values are presented near each node as SH-aLRT support (%) / ultrafast bootstrap support (%). Branch lengths were optimized by maximum likelihood on original alignment, scale bar represents substitutions/site. The colored lines correspond to Scarabaeidae subfamilies.

## Discussion

The complete circular mitogenome sequence of *O. rhinoceros* (20,898 bp) is among the largest reported in Coleoptera, and is similar in size to another scarab beetle, *Protaetia brevitarsis* (20,319 bp) (Kim et al. 2014). Mitogenome size is driven by the large non-coding (control) region that is 6,204 bp- and 5,654 bp-long in *O. rhinoceros and P. brevitarsis*, respectively. In *Popillia mutans*, another scarab beetle with a complete mitogenome sequence, the reported length of this region is only 1,497 bp (N. Song and Zhang 2018). We only recovered 17,665 bp in our Illumina-only assembly, however, and it is likely that many other beetles have larger mitogenomes, but the repetitive control regions cannot be accessed by short-read data alone. The 3,233 bp of extra control region that we recovered with long-read data includes three transcriptionally active sites. This highlights the new discoveries in mitogenomics that are likely to be made with long-read sequencing technology.

Variation in the size and nucleotide composition of the *O. rhinoceros* mitochondrial control region is not unusual in insects (Zhang and Hewitt 1997), however, there could also be technical reasons for some size discrepancies among taxa. Namely, the control region often contains tandem duplications and other repetitive sequences that present a challenge for the assembly and annotation algorithms with short-read sequence data (Tørresen et al. 2019). Second, PCR-amplification is biased against genomic regions with high AT-content that is common for the non-coding sequences, leading to the low sequencing depth in these regions, which also hinders the assembly process (Oyola et al. 2012; Gan, Linton, and Austin 2019). Library preparation and/or sequencing that is based on the PCR-amplification (e.g. Illumina) can therefore lead to AT-rich regions being underrepresented in the sequence data. The mitogenome of *O. rhinoceros* and other scarab beetles is AT-rich (N. Song and Zhang 2018), and tandem repeats can be present within the control region, making the assembly process with the PCR-based short-read sequencing technology challenging for this group. For example, three out of five scarab mitogenome assemblies recently generated with the short-read technology are incomplete and lack the control region and adjacent genes (N. Song and Zhang 2018). Our approach included the library preparation with non-PCR-amplified DNA and the long-read (ONT) sequencing, enabling us to generate a fully closed circular assembly with thousands of reads spanning the entire length of the control region. The superiority of long-read sequencing technologies to capture the long repeated, AT-rich sequences has led to the discovery of remarkable interspecific variation in the length of the intergenic repeat regions in the mitogenomes of seed beetles (Chrysomelidae), that can range between 0.1-10.5 kbp (Sayadi et al. 2017).

We also found evidence of some transcriptional activity within the control region of the *O. rhinoceros* mitogenome, but to fully characterize this pattern, more transcriptome data (from different tissues, life stages etc.) would need to be tested. Transcriptional activity within the intergenic repeat regions has been detected in mitogenomes of seed beetles (Sayadi et al. 2017), suggesting that ‘mitochondrial dark matter’ could be a source of non-coding RNAs in insects.

In the CRB mitogenome, *trnQ* gene preceded *trnI* gene, and this rearrangement was supported with thousands of long reads spanning this region. The rearranged position of *trnI* and *trnQ* genes is found in almost all species of Hymenoptera (Dowton et al. 2009), and was also reported in flatbugs (Hemiptera, Aradidae) (F. Song et al. 2016). A number of other rearrangements in trna genes have been reported in Lepidoptera and Neuroptera (Cao et al. 2012; Cameron et al. 2009), and because they all occurred between the control region (CR) and *cox1*, it has been hypothesized that this might be a ‘hotspot’ region for such changes (Dowton et al. 2009).

Quality of the CRB mitogenome assembly and annotation was validated through the phylogenetic analysis of PCGs sequence variation that correctly grouped *O. rhinoceros* with another member of the Dynastinae subfamily (Figure 2), and reconstructed the previously established relationship among several scarabid lineages (N. Song and Zhang 2018). Our *in silico*-generated PCR-RFLP marker correctly matched the CRB-G haplotype marker (Marshall et al. 2017), further supporting the high quality of the mitogenome sequence and annotation.

## Conclusions

We report the circularized complete mitochondrial genome assembly for *Oryctes rhinoceros*, the major insect pest of coconut and oil palms. The long-read ONT sequencing allowed us to identify structural variation (*trnI-trnQ* rearrangement) and span the assembly across the entire 6,203 bp-long control region that contains tandem repeats and regions of transcriptional activity. This high-quality genomic resource facilitates future development of a molecular marker toolset to help with the biosecurity and management efforts against this resurgent pest. As the first complete mitogenome for the genus *Oryctes* and the subfamily Dynastinae, and among a few for the entire scarab beetle family (Scarabaeidae), it will contribute to the resolution of higher-level taxonomy and phylogeny of phytophagous scarab beetles that remain understudied despite containing many agricultural pests.

## Acknowledgements

This project was supported by the Australian Centre for International Agricultural Research funding (HORT/2016/185), the University of Queensland (UQECR2057321) and by core funds from the Mosquito Control Laboratory at QIMR Berghofer MRI.

## Authors Contribution

**Igor Filipović:** Methodology, Investigation, Data curation, Visualization, Writing-Original draft preparation.

**James P. Hereward:** Data curation, Visualization, Writing-Reviewing and Editing.

**Gordana Rašić:** Methodology, Investigation, Resources, Writing-Reviewing and Editing.

**Gregor J. Devine:** Resources, Supervision, Writing-Reviewing and Editing.

**Michael J. Furlong:** Funding acquisition, Project administration, Supervision, Writing-Reviewing and Editing.

**Kayvan Etebari:** Conceptualization, Investigation, Resources, Funding acquisition, Supervision, Writing-Reviewing and Editing.

**Supplemental Figure 1.**
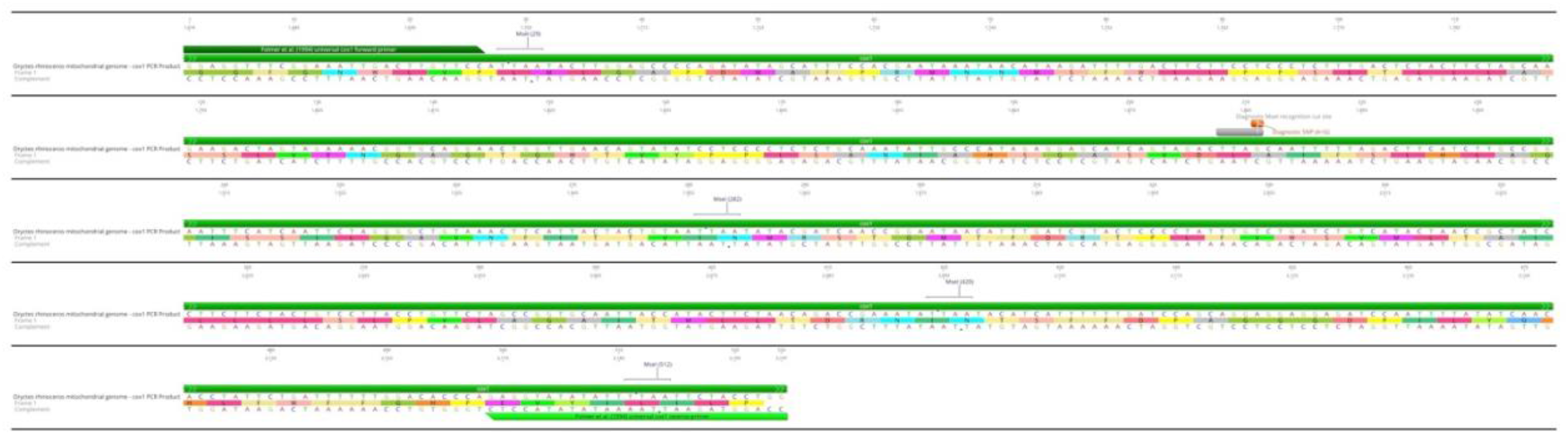
Graphical representation of *in silico* PCR-RFLP marker analysis. The amplicon from the partial cox1 gene is delineated with forward and reverse universal *cox1* primers (Folmer et al. 1994) and MseI restriction enzyme recognition sites are marked. The diagnostic SNP (Marshall et al. 2017) A>G is marked in orange. This analysis generates *in silico* fragments 253 bp, 138 bp, 92 bp, 28 bp and 13 bp-long. The fragments 253, 138 and 92 bp are visible on a 2% agarose gel in (Marshall et al. 2017) and are diagnostic for the CRB-G haplotype.

**Supplemental Figure 2.**
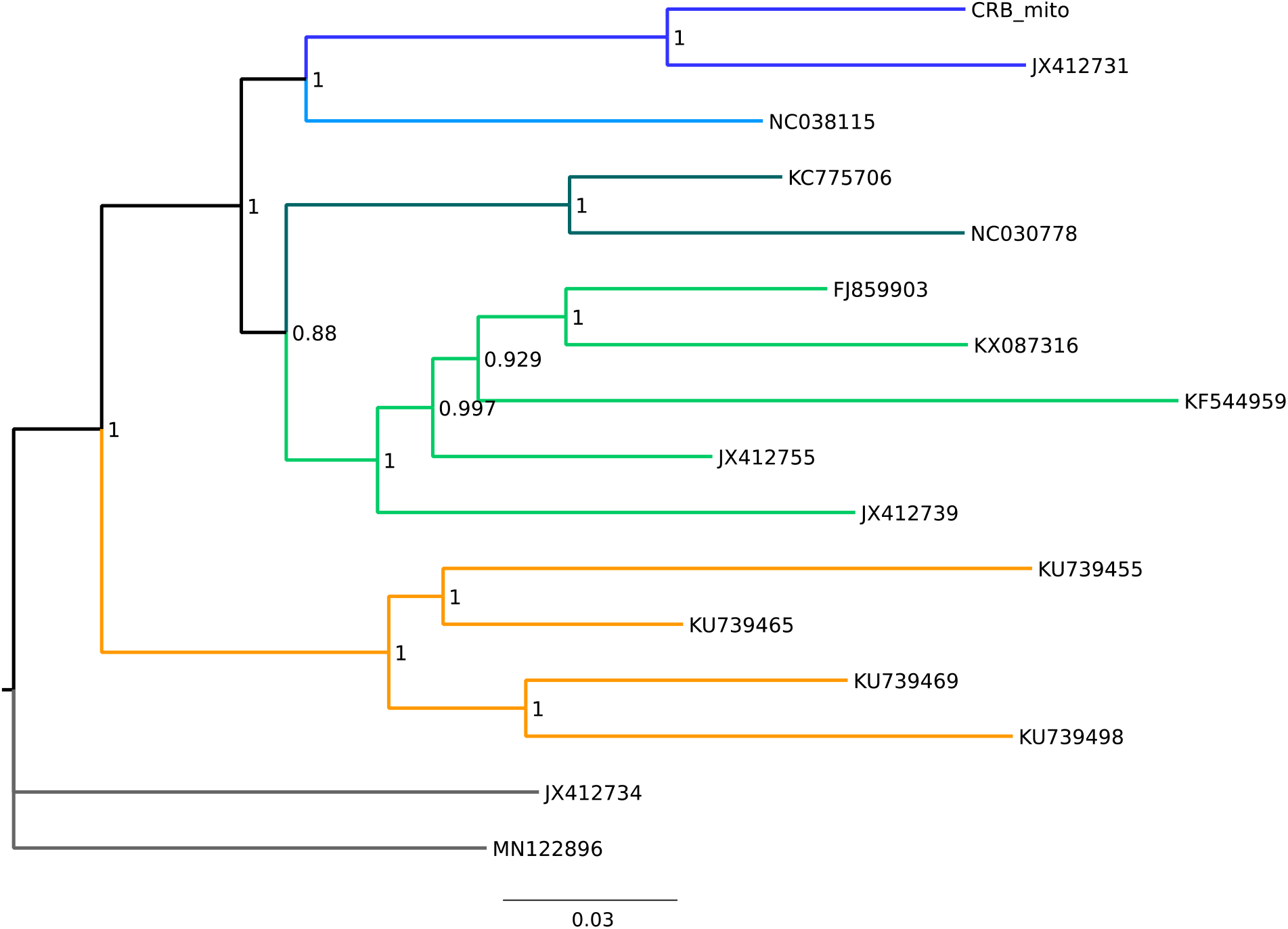
Bayesian phylogeny (MrBayes) with 13 PGSs from CRB and 15 other taxa (see Table 1). Consensus tree with branch support value as posterior probabilities (0-1).

## Notes

### Competing Interest Statement

The authors have declared no competing interest.

